# Orbitofrontal noradrenaline mediates volatility-dependent adjustment of learning rate

**DOI:** 10.64898/2025.12.11.693701

**Authors:** Hadrien Plat, Coline Chevallier, Alessandro Piccin, Alain Marchand, Jérémie Naudé, Etienne Coutureau

## Abstract

Adaptive decision-making in dynamic environments requires flexible adjustment of learning speed to balance stability and flexibility. When outcomes are highly stochastic, learners must avoid over interpreting noise and update more slowly, whereas in volatile environments where contingencies change frequently, learning should accelerate to rapidly incorporate new evidence. Theories propose that internal estimates of uncertainty tune learning rates through neuromodulatory-dependent mechanisms. Here, we investigated how noradrenergic inputs from the locus coeruleus (LC) to the orbitofrontal cortex (OFC) support adaptive learning under uncertainty. We show that rats performing a probabilistic reversal learning task exhibited behavior that was best explained by an adaptive reinforcement-learning model in which learning rates dynamically adjust according to estimated environmental volatility and stochasticity, outperforming standard fixed-rate models. Noradrenaline release in the OFC closely tracked trial-by-trial, model-derived volatility estimates around contingency changes. Disrupting LC→OFC noradrenergic inputs reproduced the model-predicted deficit in volatility-dependent adjustments of learning rate.

Together, these findings identify OFC noradrenergic signaling as a key circuit mechanism for volatility-dependent modulation of learning rates during adaptive decision-making.

## Introduction

Adaptive behavior requires anticipating the likely consequences of possible actions and update those expectations when circumstances change. When the statistical structure of the environment shifts over time, successful adaptation requires monitoring whether observed outcomes deviate from expectations and revising action–outcome (A–O) associations accordingly (1–3). Many real-world environments, however, are both stochastic (outcomes are noisy) and volatile (contingencies can change suddenly). These two forms of uncertainty place opposing demands on learning. When outcomes are highly stochastic, it becomes difficult to distinguish true changes from random fluctuations, and learning should proceed slowly to avoid overreacting to noise. In contrast, when volatility is high and contingencies shift frequently, learning should accelerate so that recent outcomes exert more influence (4–6). Financial markets offer a clear example: day-to-day price fluctuations often reflect noise, so that updating beliefs too quickly in response to these fluctuations can lead to abandoning sound investments. Yet, when broader market conditions genuinely change, maintaining outdated expectations can also be costly. The challenge is determining when variability signals a true shift rather than background noise. Thus, adaptive behavior requires dynamically balancing stability and flexibility as a function of environmental uncertainty.

Classical reinforcement-learning (RL) models typically use a fixed learning rate that determines how strongly new outcomes influence value estimates (7). In contrast, recent adaptive or meta-RL frameworks propose that the learning rate itself is dynamically regulated according to internal estimates of stochasticity and volatility, decreasing when outcomes are noisy and increasing when evidence suggests that contingencies have changed (1, 3, 8–11). These models provide a principled account of how learning speed could adjust to the demands of a dynamic environment. However, the neural mechanisms through which such volatility- and stochasticity-dependent learning-rate modulation are implemented remain unclear.

Neuromodulatory systems are well positioned to support meta-learning – dynamically adjusting the learning rate – through their widespread projections to cortical and subcortical regions (10–13). Among them, the locus coeruleus–noradrenaline (LC-NA) system has been proposed to signal unexpected events and shifts in task state (14), thereby facilitating behavioral adaptation when environmental contingencies change (12, 15–18). Recent work suggests that LC activity correlates with learning rates and uncertainty estimates in humans

(6, 19, 20) and with flexibility in rodents (21). Pharmacological studies in humans further suggest that noradrenergic signaling may modulate how strongly new outcomes update beliefs in volatile environments.

The LC is the primary source of NA in the forebrain and provides dense innervation of prefrontal regions, including the orbitofrontal cortex (OFC) (21–25). The OFC has been critically involved in updating A–O associations to support flexible, goal-directed behavior (26–29). These anatomical and functional properties position the LC→OFC pathway as a strong candidate to mediate adaptive learning under uncertainty, which we examined using a probabilistic reversal learning task that captures the demands of learning in uncertain and changing conditions.

To address this question, we combined computational modeling, fiber photometry recordings, and pathway-specific chemogenetic manipulations in rats to dissect the contribution of OFC noradrenergic signaling to flexible learning. Chemogenetic inhibition of OFC neurons selectively impaired post-reversal adaptation and reduced estimated learning rate. By varying the level of outcome stochasticity, we found that rats’ behavior was best captured by an adaptive RL model that dynamically adjusts learning rates based on estimates of volatility and stochasticity, unlike fixed-rate RL models. Fiber photometry revealed that NA release in the OFC around reversals closely tracked the trial-by-trial volatility estimates generated by the adaptive learning rate model. We then simulated a disruption of the link between volatility and learning rate to model the chemogenetic inhibition of LC→OFC projections. Selective inhibition of LC→OFC projections produced the behavioral effects predicted by disrupting volatility-driven modulation of learning rates in the model. Deficits indeed emerged specifically after reversals in the probabilistic conditions, whereas performance under deterministic contingencies remained intact.

Together, these results suggest that noradrenergic signaling in the OFC contributes to behavioral flexibility through the volatility-dependent adjustment of learning rates.

## Results

### Rats learn a history-dependent strategy in a probabilistic reversal task

To investigate how rats use reward information to guide decisions under uncertainty, we implemented a two-armed bandit probabilistic reversal learning (PRL) task (Fig. 1A). On each trial, rats chose between two levers associated with complementary reward probabilities (p(high)=1−p(low)). Whenever the high-probability lever was selected on eight consecutive trials (independently of reward delivery), reward contingencies reversed, prompting animals to adjust their choices in response to changing outcomes (Fig 1B).

**Figure 1:**
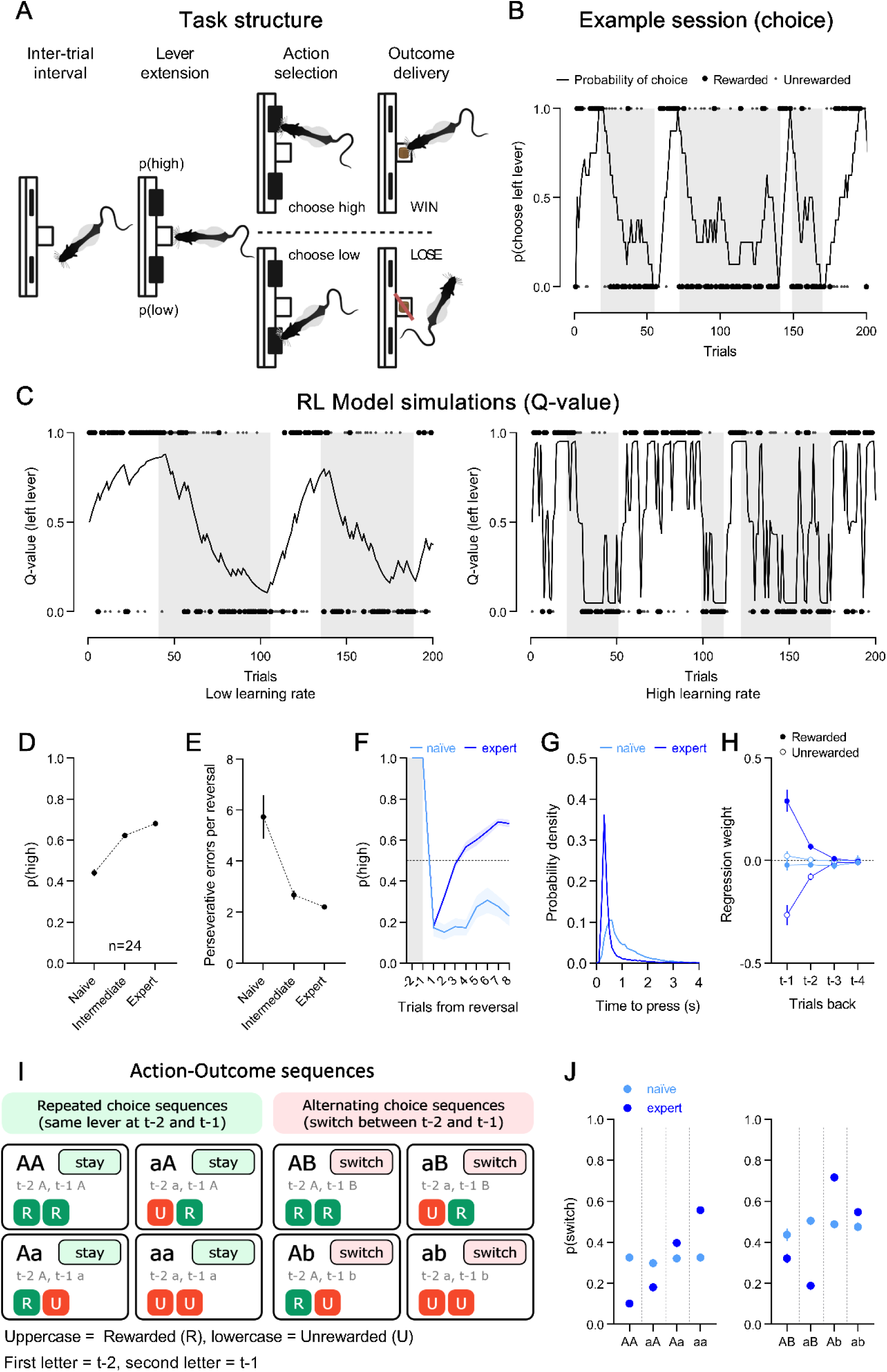
Probabilistic reversal learning task and training. (A) Task structure. Schematic representing a two-armed bandit probabilistic reversal learning task in which rats select freely between two levers yielding anti-correlated reward probabilities. Reward contingencies reversed each time the currently high-reward probability lever was selected on eight consecutive trials, requiring animals to adapt their choice strategy. (B) Example behavioral session from an expert rat. The solid line shows the rolling probability of choosing the left lever computed in an 8-trial sliding window. Large black dots indicate rewarded choices, and small grey dots indicate unrewarded choices. Rats exhibit rapid adaptation at each contingency reversal (grey shaded areas). (C) Estimated Q-value for the left lever over time in simulations of a standard reinforcement-learning model with either a low (left panel) or high (right panel) learning rate (α). The solid line shows the evolving Q-value estimate. Large black dots represent rewarded outcomes; small grey dots represent unrewarded outcomes. A low α produces slow adjustment of lever values across reversals, whereas a high α yields rapid but unstable value estimates driven by outcome noise. (D) Learning performance across training, expressed as the proportion of high-reward probability lever choices. (E) Average number of consecutive errors following reversals of contingencies (perseverative errors) across training stages. (F) Post-reversal performance, showing the proportion of high-reward probability lever-presses in the first eight trials following a contingency reversal for naïve and expert rats. (G) Frequency distribution of lever-press latencies in naïve and expert animals. (H) Logistic-regression coefficients estimating the influence of past rewarded and unrewarded trials on current decisions. Positive weight coefficient values indicate a tendency to repeat the previous choice (stay), whereas negative values indicate increased likelihood of switching. Expert in blue. Naïve in black. (I) Coding scheme used to categorize trial sequences based on lever choice and outcome history over the two preceding trials (t–2 and t–1). Upper-case letters denote rewarded outcomes and lower-case letters denote unrewarded outcomes; identical letters indicate that the same lever was chosen on trials t–2 and t–1 (stay), whereas different letters indicate a switch between these two trials. (J) Probability of switching as a function of recent action–outcome sequences as described in (H), comparing naïve and expert rats. Codes represent the combination of outcomes on trials t–2 and t–1 and the choice transition between them; same vs different letters correspond to stay *vs.* switch, respectively. Data are presented as mean ± SEM.

This exposes animals to two forms of uncertainty: volatility (due to changing reward contingencies) and stochasticity (due to probabilistic outcomes). Hence, rats are faced with the challenge of detecting meaningful changes in reward contingencies (volatility) while ignoring random noise (due to stochasticity). In a reinforcement-learning (RL) framework, this trade-off requires updating value estimates at a rate that avoids overreacting to noise while remaining responsive to meaningful change. Expressed in a standard RL Q-learning model, a rat with a low learning rate α would update the values of levers too slowly, leading to delayed adaptation after actual reversals, whereas with a high α, behavior would become overly sensitive to noise, producing erratic switching behavior (Fig. 1C). This simple model prediction suggests that rats would benefit from adjusting their learning rate depending on environmental statistics.

We examined this possibility by first training rats on 80/20 probabilistic contingency (p(high)=0.8, p(low)=0.2). Task performance improved across training, from naïve to intermediate and finally expert stages, reflected by a progressive increase in the proportion of choices to the high-probability lever (Friedman test: χ²(2)=44.33, p<0.0001; Fig. 1D). This improvement was accompanied by fewer consecutive errors following reversals (Friedman test: χ²(2)=20.58, p<0.0001; Fig. 1E) and by faster post-reversal adaptation (two-way RM ANOVA: Training stage, F(1,23)=150.2, p<0.0001; Fig. 1F), indicating that with training, rats became more proficient at extracting relevant information from noisy feedback in a volatile environment.

Rats showed no side bias (mean=0.499 ± 0.006 SEM) and responded quickly (latency=0.672 ± 0.062 s), with reaction times influenced by training stage (Fig. 1G). To quantify how past events shaped choice behavior, we next applied a logistic regression predicting stay/switch decisions from reward outcomes over recent trials (Fig. 1H; see Methods). Positive weight regression coefficients indicate a tendency to repeat the previous choice, whereas negative coefficients indicate switching. Expert rats displayed strong positive weights following rewarded trials and strong negative weights following unrewarded trials, demonstrating a sensitivity to feedback across multiple past trials. In contrast, naïve rats showed no significant influence of past outcomes on their decisions.

To disentangle the respective influence of past outcomes and past choices on switching, we computed the conditional probability of switching for all possible two-trial choice–outcome sequences preceding the current decision (Fig. 1I). Naïve rats increased switching primarily when they had switched on the previous trial, independent of whether the past outcomes were rewarded or unrewarded (three-way RM ANOVA: Action t-1, F(1,23)=73.93, p<0.0001; Outcome t-1, F(1,23)=0.4672, p=0.501; Outcome t-2, F(1,23)=0.5010, p=0.486; three-way interaction, F(1,23)=6.41, p=0.0186; Fig. 1J). In contrast, expert rats used both recent choice and outcome history to guide switching (Fig. 1J), with significant main effects and an interaction between these factors (three-way RM ANOVA: Action t-1, F(1,23)=128.8, p<0.0001; Outcome t-1, F(1,23)=956.3, p<0.0001; Outcome t-2, F(1,23)=5.997, p=0.0224; three-way interaction, F(1,23)=9.553, p=0.0052). This emergence of structured, history-dependent switching is consistent with RL principles and indicates that rats adapted to the demands of the task by learning to use recent feedback to guide behavior.

### Inhibiting OFC neurons disrupts learning rate during probabilistic reversal learning

To test whether the orbitofrontal cortex (OFC) contributes to adaptive learning in uncertain environments, we chemogenetically inhibited CaMKII+ neurons in the ventral and lateral OFC (VO/LO) using an AAV8-CaMKIIα-hM4Di-mCherry construct (Fig. 2A–B). After training on the 80/20 schedule, rats were tested under vehicle (VEH) and DCZ inhibition conditions (see Methods).

**Figure 2:**
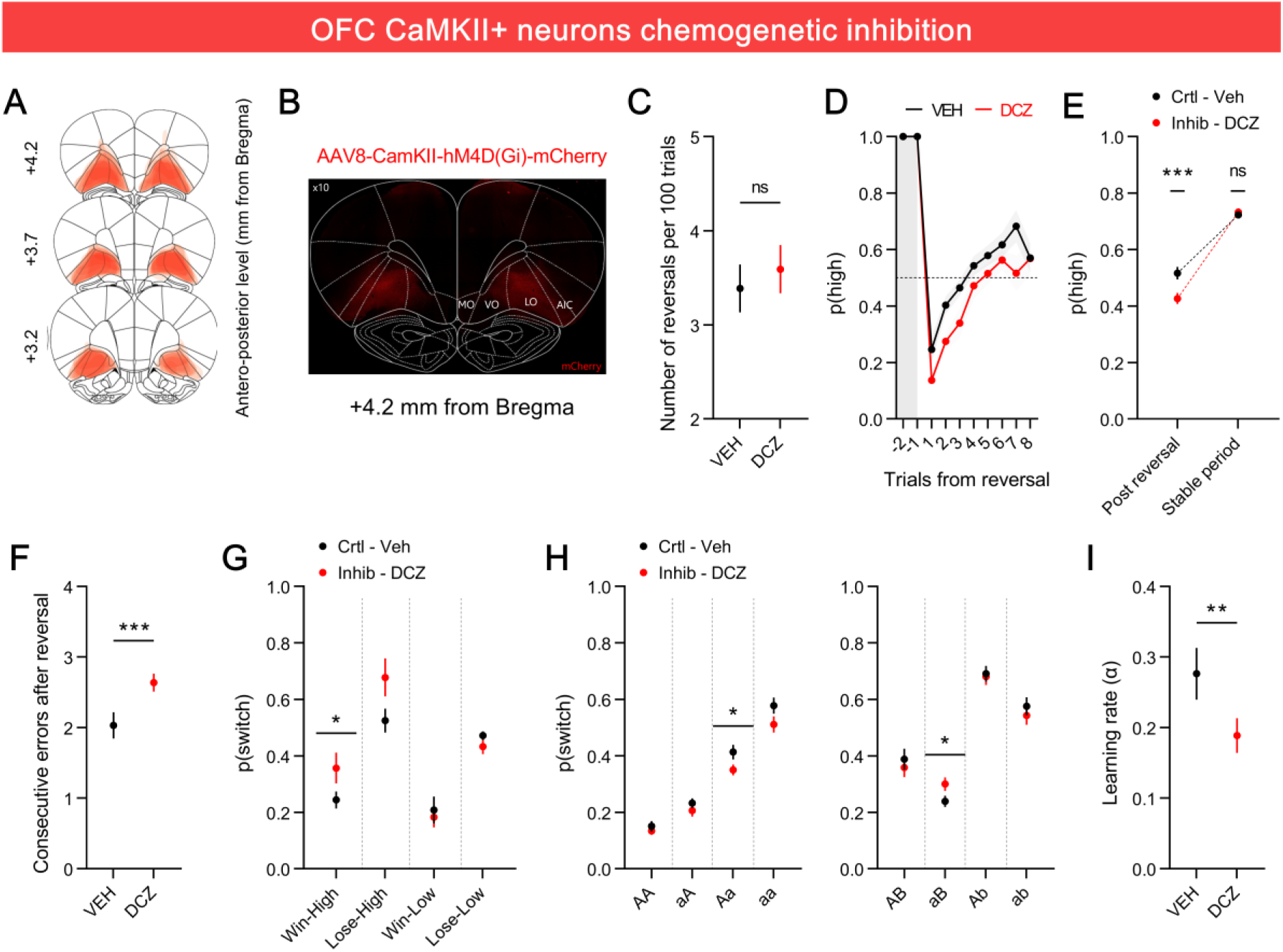
Inhibiting OFC neurons disrupts flexible adaptation during probabilistic reversal learning. (A) Schematic representation of viral expression across subjects, showing the extent of AAV8-CaMKII-hM4D(Gi)-mCherry expression within the orbitofrontal cortex (OFC; VO and LO subregions). Each animal is represented as a separate layer (n = 12). (B) Representative fluorescence image illustrating AAV8-CaMKII-hM4D(Gi)-mCherry expression within the OFC (+4.2 mm from Bregma). (C) Mean number of reversals completed per 100 trials under vehicle (VEH) and deschloroclozapine (DCZ, 0.1 mg/kg, i.p.) treatments. (D) Post-reversal adaptation showing the trial-by-trial probability of selecting the high-probability lever following contingency reversals under VEH and DCZ conditions. (E) Proportions of high-reward probability lever-presses during the post-reversal period (left) and during stable phases (right) under VEH and DCZ treatments (***p < 0.001; n.s., not significant). (F) Average number of consecutive (perseverative) errors following reversals under VEH and DCZ conditions (***p < 0.001). (G) Probability of switching levers following different outcome types: rewarded (win), unrewarded (lose), high-probability (high), or low-probability (low) lever presses in the post-reversal phase (*p < 0.05). (H) Conditional probability of switching based on recent action–outcome sequences across the last two trials under VEH and DCZ treatments (*p < 0.05). (I) Learning rates (α) estimated from a standard reinforcement-learning model fitted to behavioral data under VEH and DCZ conditions (**p < 0.01). Data are presented as mean ± SEM.

While DCZ treatment did not affect the total number of reversals completed (paired t-test: t(11)=0.6606, p=0.5224; Fig. 2C), silencing OFC neurons significantly decreased rats’ ability to adjust to new contingencies after reversals, while leaving their performance during stable periods unaffected (two-way RM ANOVA: treatment × phase, F(1,11)=17.00, p=0.0017; post-hoc: post-reversal, p=0.0005; stable, p=0.83; Fig. 2D–E). This impairment was marked by a significant increase in consecutive errors following contingency reversals in DCZ treated sessions (paired t-test: t(11)=4.52, p=0.0009; Fig. 2F), indicating increased perseverative behavior. In addition to initial perseveration on the previously highly-rewarded lever, rats also showed increased switching away from the new high-probability lever during the post-reversal period (paired t-test: t(11)=2.826, p=0.0165; Fig. 2G), suggesting impaired updating following reversals.

Conditional switching probability-based analyses revealed that OFC inhibition also selectively altered how rats used past choice-outcome information (Fig. 2H). Specifically, rats were less likely to switch after experiencing an Aa sequence (paired t-test: t(11)=2.451, p=0.0322) and more likely to switch following aB sequences (paired Wilcoxon test: p=0.0093). Overall, these results suggest that OFC activity plays a key role in integrating prior action-outcome information to guide decision-making in an uncertain environment. To further understand how OFC inhibition affects learning dynamics, we fitted a standard RL model to the behavioral data. OFC inhibition significantly reduced apparent learning rates, confirming diminished sensitivity to recent outcomes when updating value estimates (paired t-test: t(11)=3.15, p=0.0093; Fig. 2I). Overall, these results show that inhibiting OFC neurons disrupts flexible learning by impairing outcome integration and reducing apparent learning rates, leading to altered post-reversal adaptation.

### Rats adaptively adjust learning rates based on volatility and stochasticity estimates

To further determine how the OFC contributes to learning regulation, we next investigated how rats adjust their learning rates as a function of environmental stochasticity (noise) and volatility (frequency of change). We based our approach on the assumption that changes in reward should induce faster adaptation to contingency shifts when noise is low. In other words, outcome feedback is more informative in deterministic environments and becomes progressively less informative about environmental changes (volatility) as stochasticity increases. We formalized this hypothesis in the RL model, showing that there is an optimal learning rate α that maximizes reward in the 80/20 PRL task (Fig. 3B). This optimal α increased with reward predictability, with less stochastic environments favoring higher optimal learning rates (Fig 3C). The model thus suggests that rats’ learning rates would benefit from adjusting to the statistical structure of the environment, enabling efficient adaptation to genuine change while minimizing overreactions to noise.

**Figure 3:**
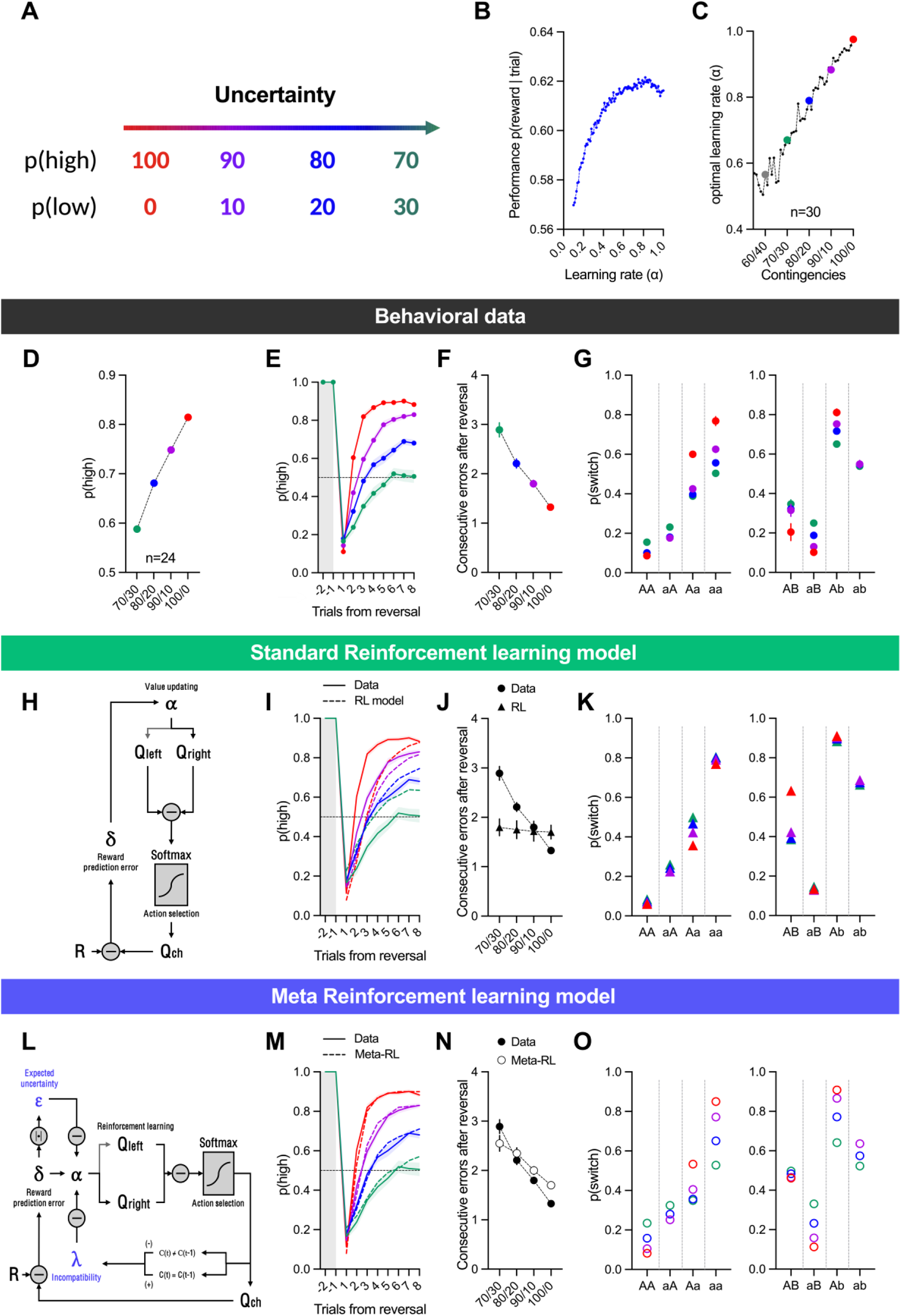
Adaptive learning in rats is best explained by a meta-reinforcement learning model. (A) Diagram illustrating the set of contingencies used during behavioral testing: 70/30 in green, 80/20 in blue, 90/10 in purple, 100/0 in red, representing a gradient of environmental uncertainty from high (stochastic) to low (deterministic). (B) Simulation results from a standard reinforcement-learning (RL) model (n = 30 agents) illustrating the impact of increasing learning rates (α) on performance under 80/20 contingencies. (C) Optimal learning rates derived from the standard RL model as a function of reward stochasticity across contingencies. (D) Behavioral performance across contingencies, showing the overall proportion of high-reward probability lever-presses (n = 24). (E) Trial-by-trial performance aligned to contingency reversals. Curves represent the proportion of high-reward probability lever-presses in the eight trials following each reversal across contingencies. (F) Average number of consecutive errors after reversals as a function of contingency, showing slower adaptation in more stochastic conditions. (G) Conditional switching probabilities based on recent action–outcome sequences across contingencies. Left, sequences in which the same lever was chosen on the two previous trials (AA, aA, Aa, aa). Right, sequences in which different levers were chosen (AB, aB, Ab, ab). (H) Schematic of the standard Q-learning model, in which action values (Q) are updated from reward prediction errors (δ) weighted by a fixed learning rate (α). (I–K) Predictions from the standard RL model: (I) post-reversal performance, (J) consecutive errors, and (K) conditional switching probabilities. The standard RL model fails to capture key features of behavior, such as adaptive switching after unrewarded trials (L) Schematic of the Meta-Reinforcement Learning (Meta-RL) model, in which the learning rate (α) is dynamically modulated by internal estimates of stochasticity (ε) and environmental volatility (λ). (M–O) Predictions from the Meta-RL model: (M) post-reversal adaptation, (N) consecutive errors, and (O) conditional switching probabilities across contingencies. The adaptive Meta-RL model reproduces the observed behavioral flexibility across stochastic environments. Data are presented as mean ± SEM.

To test this prediction, we manipulated outcome stochasticity across different reward schedules (70/30, 80/20, 90/10, and 100/0) in the same rats as in Experiment 1 (n=24), across separate sessions (Fig. 3A; see Methods), to examine whether learning rates were modulated by stochasticity in accordance with the theoretical optimum predicted by the model. Rats’ performance improved as stochasticity decreased, reflected by a higher likelihood of selecting the advantageous lever (Friedman test: χ²(3)=72.00, p<0.0001; Fig. 3D). Following reversals, rats adapted more rapidly in low-stochasticity environments, switching faster toward the newly rewarded option (two-way RM ANOVA: main effect of Contingency, F(3,69)=250.0, p<0.0001; Trials × Contingency interaction, F(21,483)=20.41, p<0.0001; Fig. 3E). Moreover, perseveration decreased as stochasticity declined, with fewer consecutive errors after reversals in more deterministic schedules (Friedman test: χ²(3)=61.04, p<0.0001; Fig. 3F), consistent with the idea that noisier environments require greater evidence accumulation before adaptation onset.

To assess how rats adjusted their choices based on recent experience, we analyzed switching behavior using the trial-sequence coding system shown in Fig. 1H (Fig. 3G). Rats consistently switched more often following unrewarded outcomes than rewarded ones across all contingencies (three-way RM ANOVA: Outcome t−1, F(1,23)=3377, p<0.0001; Outcome t−2, F(1,23)=97.48, p<0.0001; Contingency, F(3,69)=48.54, p<0.0001; Contingency × Outcome t−1, F(3,69)=218.2, p<0.0001; all other interactions p>0.07; Fig. 3G). Switching also depended on the structure of recent action–outcome (A–O) sequences. When rats repeated the same lever on the previous two trials (AA, aA, Aa, aa), switching increased with both the number and recency of losses (three-way RM ANOVA: Outcome t−1, F(1,23)=1534, p<0.0001; Outcome t−2, F(1,23)=275.8, p<0.0001; Contingency × Outcome t−1, F(2,46)=40.97, p<0.0001; Fig. 3G, left). When rats had just switched levers, switching again depended on recent outcomes (three-way RM ANOVA: Outcome t−1, F(1,23)=1001, p<0.0001; Outcome t−2, F(1,23)=279.1, p<0.0001; Fig. 3G, right), and stochasticity again modulated how feedback from recent trials guided behavioral updating (Contingency × Outcome t−1, F(2,46)=12.25, p<0.0001; Contingency × Outcome t−2, F(2,46)=8.05, p<0.01).

Importantly, outcome stochasticity further modulated these patterns. For repeated-choice sequences (left panel), switching after a *rewarded* trial was highest in the most stochastic schedule (70/30), as confirmed by significant effects of contingency for both AA and aA sequences (one-way RM ANOVA: F(3,69)=33.03, p<0.0001; F(2,46)=7.486, p<0.01). Post hoc tests showed that switching in the 70/30 condition was greater than in all other contingencies (all p<0.01). In contrast, switching after a single *unrewarded* trial (Aa) was greatest in the deterministic schedule (100/0) (one-way RM ANOVA: F(3, 69)=71.56, p<0.0001; post-hoc: 100/0 > 70/30, 80/20, 90/10, all p<0.0001), where each negative outcome provides strong evidence for a contingency change. Conversely, in stochastic environments an unrewarded outcome is ambiguous, as it may reflect either a true reversal or simply noise in reward delivery. Following *two unrewarded* trials (aa), switching also increased as stochasticity decreased (Friedman test: χ²(3)=51.35, p<0.0001), indicating that rats accumulated more evidence before changing choices in noisier environments. For alternating-choice sequences (right panel), switching remained stable for *uninformative* outcome pairs (AB: Friedman test: χ²(3)=8.584, p=0.0354; post-hoc 100/0 < 70/30, p=0.0370; ab: F(2, 46)=0.096, p=0.9084.). In contrast, switching varied strongly with stochasticity for *informative* opposite-outcome combinations (aB: F(3,69)=57.18, p<0.0001; Ab: Friedman test: χ²(3)=38.15, p<0.0001).

To test whether a fixed learning rate could account for rats’ behavior, we simulated performance across stochasticity levels using a standard Q-learning model with a constant α (Fig. 3H). This model failed to reproduce key features of the data. First, it generated only minor differences in post-reversal performance across stochasticity conditions (Fig. 3I) and produced an almost constant number of consecutive perseverative errors across conditions (Fig. 3J), in stark contrast to the behavioral results.

Looking at switching as a function of recent A–O sequences, the failures of the fixed-α model fell into two clear categories (Fig. 3K). (1) Inverse predictions: for sequences in which rats lost once or twice on the same lever (Aa, aa) or won on different levers (AB), the model predicted more switching in noisy environments, whereas rats showed the opposite pattern, switching more in deterministic environments, where outcomes are more informative. (2) Lack of stochasticity effects: for sequences with conflicting outcomes across levers (aB, Ab), rats’ switching behavior scaled with outcome stochasticity, but the model predicted identical switching probabilities across contingencies, failing to capture this graded adaptation. Together, these discrepancies demonstrate that a constant-learning-rate model fails to account for the stochasticity-dependent behavioral adaptations observed in rats, falsifying the fixed-α hypothesis and suggesting the need for an alternative model capable of accommodating such flexibility.

To overcome the limitations of fixed learning-rate models, we developed a meta-RL model with an adaptive learning rate, in which α dynamically adjusts based on internal estimates of outcome stochasticity (ε) and environmental volatility (λ) (Fig. 3L). In this framework, unsigned prediction errors (PEs) provide an index of expected uncertainty and are used to estimate ε (stochasticity), while volatility (λ) is estimated from A–O relationships that deviate from expectations, what we term “incompatibility” (see Methods). When an animal maintains the *same* choice strategy but receives worse-than-expected outcomes, the model interprets this as a rise in volatility, increasing α to accelerate updating. Likewise, if a *switch* yields better-than-expected outcomes, λ also increases, again elevating α. Conversely, large unsigned PEs (both negative and positive) occurring regardless of choice are interpreted as high stochasticity (high ε), leading the model to decrease α to avoid overreacting to noise. Thus, with only simple computations derived from PEs, the adaptive learning-rate model provides a principled mechanism for modulating learning in response to the combined demands of volatility and stochasticity.

The adaptive learning-rate model successfully captured the behavioral effects of outcome stochasticity on reversal performance. It reproduced the differences in post-reversal choice accuracy across contingency schedules (Fig. 3M) and, unlike the fixed-α model, correctly predicted reduced perseveration when outcomes were more reliable (Fig. 3N). These findings support the idea that rats dynamically adjust their learning rate based on environmental structure, updating rapidly when outcomes are reliable and adjusting more cautiously when they are noisy.

Finally, the adaptive model also accurately reproduced the conditional switching patterns observed in rats (Fig. 3O). It captured both (i) the hierarchical structure of switching based on repeated versus alternating A–O sequences (AA < aA < Aa < aa and aB < AB < ab< Ab) and (ii) how switching scaled with stochasticity for each sequence. By contrast, the standard RL model either predicted the wrong direction of effects (Aa, aa, AB) or failed to produce stochasticity-dependent modulation (aB, Ab). Together, these results show that incorporating volatility- and stochasticity-dependent adjustments to α enables an RL model to replicate key features of adaptive choice behavior observed in animals.

### Orbitofrontal noradrenergic release reflects estimated volatility

We next sought a neurobiological mechanism that could support the adaptive modulation of learning rates in the OFC (Fig. 2–3). Because the locus coeruleus–noradrenaline (LC–NA) system is known to signal unexpected changes in the environment and promote behavioral flexibility, we asked whether NA release in the OFC might reflect computations related to stochasticity and volatility. To address this, we monitored NA dynamics in the OFC using GRAB-NE1m fiber photometry (Fig. 4A–B). Animals were trained to expert performance on the 80/20 schedule prior to recordings, showing comparable behavior to previous experiments (Fig. Supp. 1A–I).

**Figure 4:**
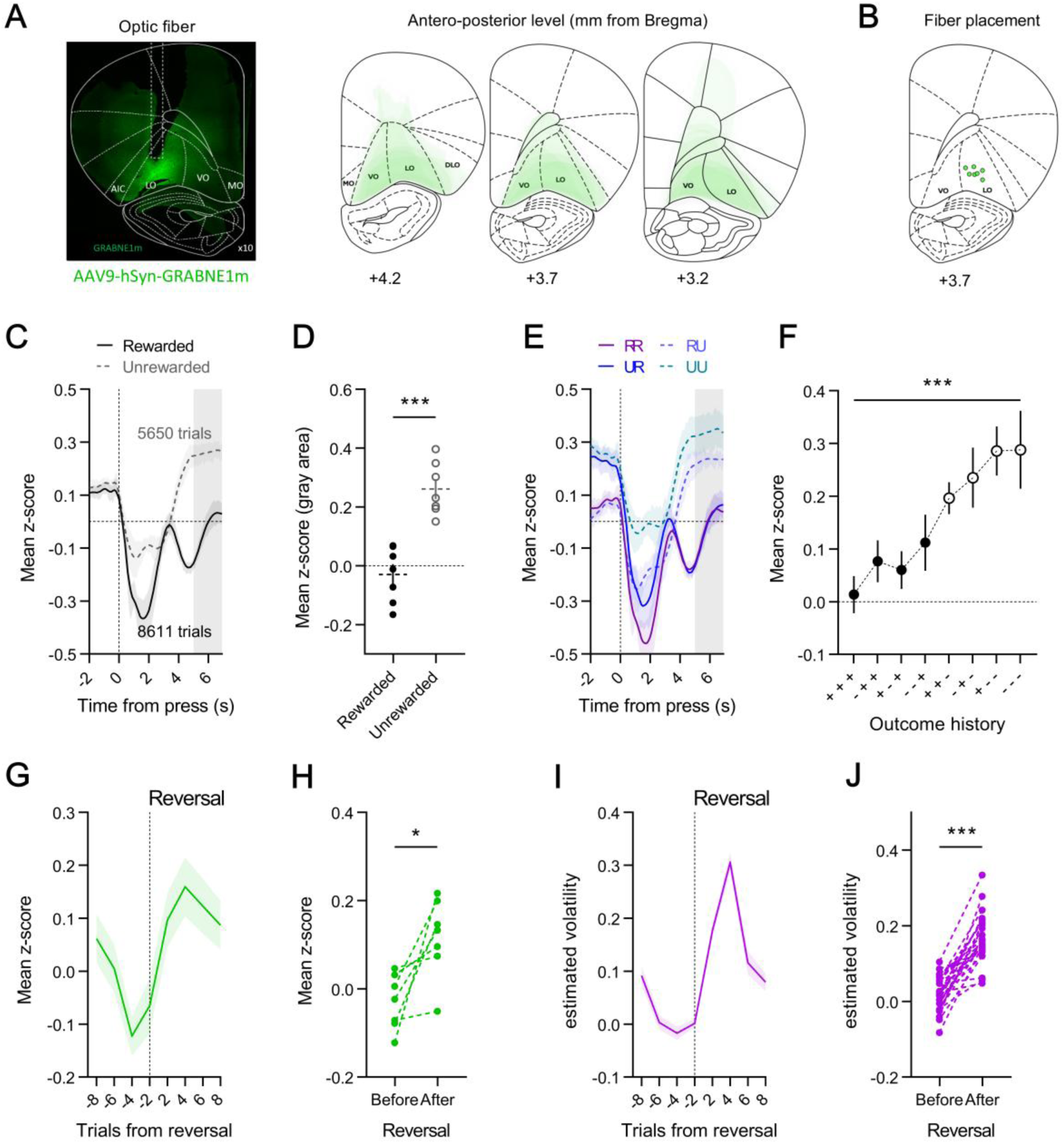
Orbitofrontal noradrenaline signals reflect model-estimated volatility. (A) Expression and spread of AAV9-hSyn-GRABNE1m in the orbitofrontal cortex (OFC). Left, representative fluorescence image showing optic-fiber placement; right, reconstructed injection sites across antero-posterior levels from Bregma (+4.2 to +3.2 mm). n = 7 rats. (B) Map of unilateral optic-fiber placements used for OFC fiber-photometry recordings. (C) Trial press-aligned NA dynamics (z-scored fluorescence) for rewarded (black) and unrewarded (gray) outcomes. Shaded region indicates the analysis window used in (D). (D) Mean NA signal in the post-outcome epoch for rewarded versus unrewarded trials. (***p < 0.001). (E) NA dynamics as a function of the combinations of current and previous trial outcomes: RR (rewarded–rewarded), RU (rewarded–unrewarded), UR (unrewarded–rewarded), and UU (unrewarded–unrewarded). (F) Pre-press NA activity as a function of detailed outcome history across the three preceding trials (t₋₃, t₋₂, t₋₁), with “+” indicating rewarded and “–” indicating unrewarded outcomes (***p < 0.001). (G) Mean NA activity aligned to contingency reversals (over eight trials before and after the switch). NA levels declined across stable pre-reversal trials and transiently increased immediately after reversals. (H) Comparison of mean NA activity before and after reversal, showing significantly higher post-reversal levels (*p < 0.05). (I) Trial-by-trial estimates of volatility from the adaptive learning-rate model aligned to reversals, displaying a temporal profile closely matching OFC NA dynamics. (J) Mean estimated volatility before vs. after reversal, showing a significant increase (***p < 0.001). Data are presented as mean ± SEM.

We first examined whether OFC–NA responses reflected immediate trial outcomes (t₀). NA release showed a biphasic, outcome-dependent pattern across the trial (Fig. 4C). NA decreased at lever press for both rewarded and unrewarded trials (two-way RM ANOVA: Phase, F(2,12)=19.00, p=0.0002; Outcome, F(1,6)=69.70, p=0.0002; Phase × Outcome interaction, F(2,12)=21.55, p=0.0001), with a larger decrease following rewarded outcomes (post-hoc, p=0.0003). Following unrewarded outcomes, NA activity rose over the inter-trial interval (post-hoc, p=0.0205), whereas after rewarded outcomes, NA continued to decline (post-hoc, p=0.0047), resulting in lower NA levels at the end of the trial (paired t-test: t(6)=6.719, p=0.0005; Fig. 4D).

We next asked whether NA also reflected recent outcome history. Pre-press NA levels were higher following an unrewarded trial on t₋₁ (two-way RM ANOVA: Outcome t₋₁, F(1,6)=24.34, p=0.0026; Fig. 4E), indicating that NA carried information from the previous choice. To determine whether this influence extended further back in time, we grouped trials according to the pattern of outcomes across the three preceding trials (t₋₃, t₋₂, t₋₁). Pre-press NA activity differed across outcome histories (one-way RM ANOVA: F(7,42)=6.422, p<0.0001; Fig. 4F), suggesting that NA release is sensitive to the pattern of recent outcomes over multiple trials. Together, these results indicate that OFC–NA responses reflect both current feedback and a graded influence of recent outcome history. We then examined how OFC–NA activity responded to changes in environmental contingencies. To address this, we aligned NA release to reversal events and quantified its dynamics over a sixteen-trial window around the switch (Fig. 4G). NA decreased across pre-reversal trials, when animals were stably exploiting the advantageous option, then exhibited a clear transient increase immediately after reversals (one-way RM ANOVA: F(7,42)=2.957, p=0.0130; Fig. 4G), before returning toward baseline as behavior adapted. Across animals, post-reversal NA was consistently higher than pre-reversal NA (paired t-test: t(6)=3.320, p=0.0160; Fig. 4H), indicating that OFC–NA activity increases when outcomes contradict established expectations.

We next asked whether the OFC–NA signal could reflect internal representations of environmental uncertainty. To address this, we compared OFC–NA dynamics to the trial-by-trial volatility estimates generated by the adaptive learning-rate model. In the model, estimated volatility exhibited a strikingly similar temporal profile to OFC–NA activity, declining across stable trials preceding the reversal before sharply rising immediately after the contingency change and gradually returning to baseline (one-way RM ANOVA: F(7,203)=49.43, p<0.0001; Fig. 4I). As for OFC-NA activity, volatility was higher after reversals compared to the stable period that preceded them (paired t-test: t(29)=11.77, p<0.0001; Fig. 4J), indicating that OFC–NA may track fluctuations in environmental uncertainty over time.

Overall, these findings suggest that noradrenergic signaling in the OFC dynamically tracks unexpected changes in outcomes, integrates recent feedback over multiple trials, and closely parallels model-derived estimates of volatility. This supports a role for OFC–NA in signaling environmental change and in modulating learning adjustments when behavioral strategies must be updated.

### Chemogenetic inhibition of Locus Coeruleus → OFC neurons reproduces model-predicted effects of volatility-dependent learning-rate adjustments

If NA in the OFC contributes to volatility-dependent adjustments of the learning rate, then affecting this signal should produce specific behavioral impairments. To test this idea, we used two complementary approaches. To generate behavioral predictions, we simulated a reduction in volatility-dependent adjustments of the learning rate (α) within our adaptive Meta-RL framework (Fig. 5B). We then evaluated this prediction experimentally by chemogenetically inhibiting LC→OFC noradrenergic projections using a retrograde CAV2-PRS-hM4Di construct injected into the OFC (Fig. 5A; Fig. Supp. 2A).

**Figure 5:**
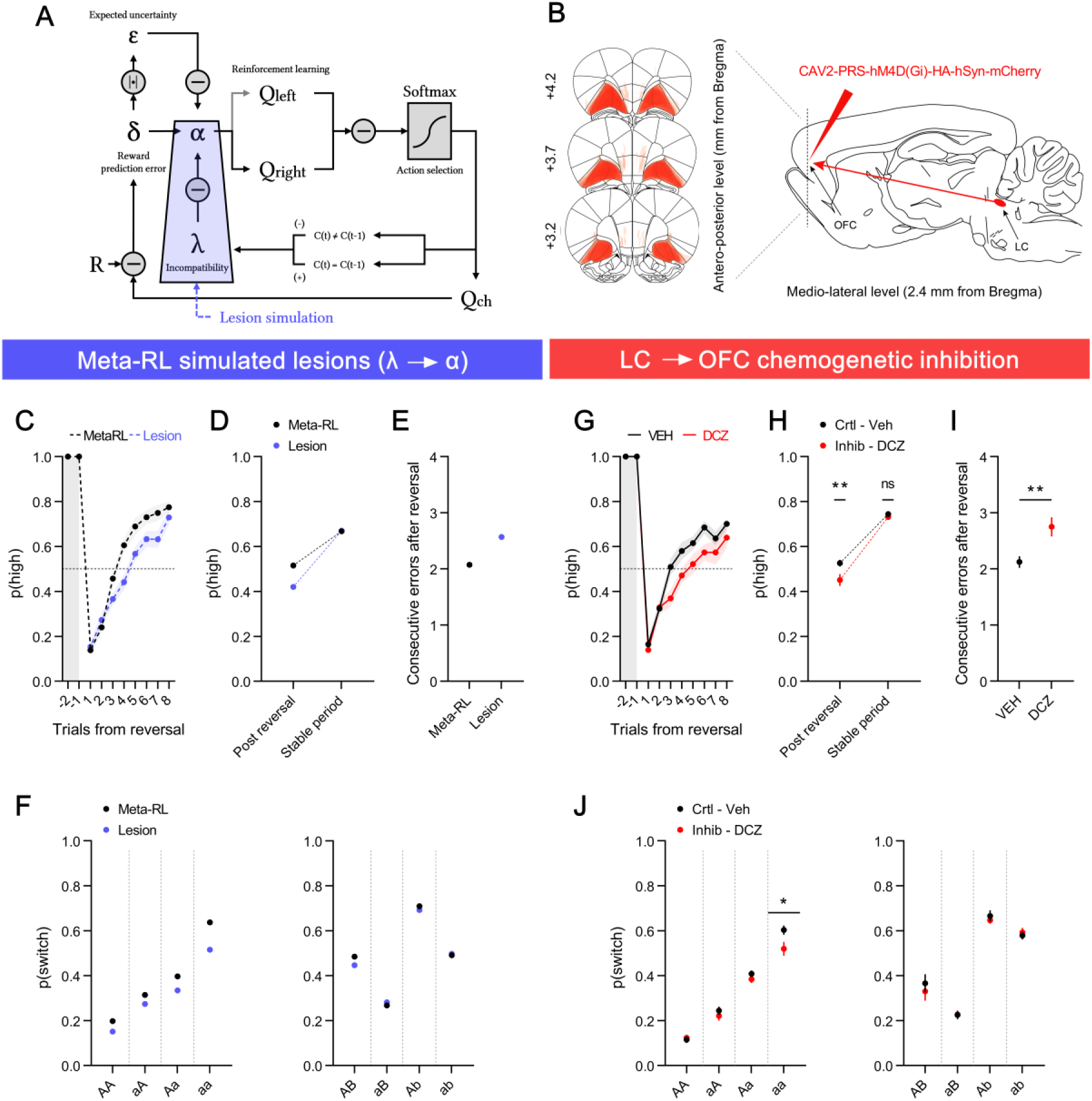
Disrupting LC→OFC noradrenergic input reproduces model-predicted impairments in adaptive learning. (A) Schematic of the Meta-Reinforcement Learning (Meta-RL) model, showing a simulated lesion of the volatility-to-learning-rate link (λ→α), used to model disruption of volatility-dependent learning-rate adjustments. (B) Anatomical schematic of the chemogenetic experiment. Left: extent of CAV2-PRS-hM4Di-HA-hSyn-mCherry expression in LC neurons projecting to the OFC, represented as stacked outlines across antero-posterior levels (+4.2 to +3.2 mm from Bregma). Right: diagram of the LC→OFC pathway and bilateral viral injection sites (n = 16 rats). (C–F) Meta-RL lesion simulations under probabilistic conditions. (C) Simulated performance across trials following reversals under intact (black) and λ→α-lesioned (blue) Meta-RL models. (D) Proportion of high-reward probability lever-presses during post-reversal versus stable periods in the same simulations. (E) Mean number of consecutive errors following reversals, showing increased perseveration in the lesioned model. (F) Simulated conditional switching probabilities based on recent action–outcome sequences (left: same lever, right: different levers), showing reduced switching after unrewarded outcomes in the lesioned model. (G–J) In vivo LC→OFC inhibition under probabilistic conditions. (G) Behavioral performance across trials following reversals under vehicle (VEH, black) and DCZ (red) treatment. (H) Proportion of high-reward probability lever-presses during post-reversal and stable periods under VEH and DCZ (treatment × phase, **p < 0.01; post-reversal **p = 0.0029; stable n.s.). (I) Mean number of consecutive errors following reversals, showing increased perseveration after LC→OFC inhibition (**p < 0.01). (J) Conditional switching probabilities under VEH and DCZ, revealing reduced switching after unrewarded outcomes (*p < 0.05). Data are presented as mean ± SEM.

In the 80/20 probabilistic condition, the computational lesion predicted a selective deficit in post-reversal adaptation, characterized by a slower increase in choosing the newly rewarded lever (Fig. 5C), while leaving performance during stable phases intact (Fig. 5D). It further predicted increased perseveration after reversals (Fig. 5E) and a reduction in switching after unrewarded outcomes when the same lever was repeated, without affecting switching following rewarded trials (Fig. 5F).

In vivo LC→OFC inhibition produced the same behavioral deficits. Performance during stable blocks was unchanged, while adaptation following reversals was significantly impaired (two-way RM ANOVA: Treatment × Session phase: F(1,15)=5.259, p=0.0367; post-hoc: post-reversal, p=0.0029; stable, p=0.7794; Fig. 5G–H), despite animals completing a similar number of reversals under both conditions (paired t-test: t(15)=0.905, p=0.3798; Fig. Supp. 2B). LC→OFC inhibition increased consecutive errors after reversals (paired t-test: t(15)=3.214, p=0.0058; Fig. 5I) and selectively reduced switching after losses on the new low-probability lever (paired t-test: t(15)=3.565, p=0.0028; Fig. Supp. 2C). Moreover, consistent with model predictions of reduced switching after unrewarded outcomes when the same lever was repeated (Fig. 5F), LC→OFC inhibition decreased switching after repeated omissions on the same lever “aa” (paired t-test: t(15)=2.803, p=0.0134; Fig. 5J). A two-way RM ANOVA on switching after unrewarded trials (ΣOmission: F(2,30)=20.16, p<0.0001; Treatment: F(1,15)=9.407, p=0.0078; ΣOmission × Treatment: F(2,30)=1.872, p=0.1714; Fig. Supp. 2D) showed that the treatment effect was most pronounced when more than one consecutive omission occurred.

We next examined whether the same manipulation would produce similar effects under deterministic conditions. In the 100/0 condition, the computational lesion predicted no behavioral impairment, with post-reversal performance, perseveration, and switching behavior remaining unchanged (Fig. Supp. 3A–D).

In vivo, LC→OFC inhibition produced the same outcome: DCZ treatment did not alter post-reversal performance (two-way RM ANOVA: Treatment, F(1,15)=0.074, p=0.7893; Treatment × Session phase interaction, F(1,15)=0.517, p=0.4831; Fig. Supp. 3E–F), perseveration (paired t-test: t(15)=0.329, p=0.7470; Fig. Supp. 3G), or conditional switching in the deterministic condition (paired t-tests, all p>0.05; Fig. Supp. 3H). Thus, the model accurately predicted both the presence of behavioral deficits in the probabilistic environment and their absence in the deterministic one.

Together, these results reveal a clear correspondence between the computational predictions and the behavioral effects of LC→OFC inhibition, reproducing the impact of disrupting volatility-dependent learning-rate adjustments.

## Discussion

Updating action-outcome associations is essential for adapting behavior in uncertain environments. Here we combined behavioral, photometry, chemogenetic, and computational approaches to identify noradrenergic signaling in ventrolateral OFC as a key mechanism supporting this adaptive process. Silencing OFC CaMKII⁺ neurons impaired post-reversal adaptation and reduced model-derived learning rates. Behavior was better captured by an adaptive reinforcement-learning (RL) model in which learning rates are adjusted trial-by-trial according to estimates of environmental volatility and stochasticity, whereas fixed-rate models failed to reproduce behavior across uncertainty levels. Fiber-photometry recordings revealed that noradrenaline (NA) release in the OFC closely tracked model-derived volatility estimates, decreasing before and increasing after reversals. Finally, LC→OFC inhibition caused selective deficits in probabilistic conditions, as predicted by disrupting volatility-dependent learning-rate control.

### Behavioral analysis and Modelling approach

A large body of work has modeled reversal behavior using classical RL frameworks with fixed parameters (30), incorporating refinements such as separate learning rates for positive and negative outcomes (29), choice-stickiness terms (31), counterfactual updating (30), or value forgetting (28, 32). Although these models achieve good fits - penalized-likelihood scores (e.g., BIC, AIC) on the whole choice series- they can underestimate the sharp adjustments that follow reversals. Furthermore, a model with fixed parameters can fit behavior under a single volatility and stochasticity level, but may fail to capture qualitative patterns of adaptation across uncertainty conditions. Following Palminteri, Wyart & Koechlin (33), we therefore evaluated our models not only by their predictive fit but also by their generative validity. When simulated, the fixed-α model failed to reproduce key behavioral patterns across varying stochasticity levels, indicating that such models lack the flexibility required to learn in dynamic settings. In environments that change over time, learners must decide how much past experience should influence current behavior, and how quickly to update when contingencies shift (34). Adaptive learning models formalize this principle. Piray and Daw (9) developed a model in which the learning rate is dynamically adjusted according to two estimates: stochasticity (random outcome noise) and volatility (genuine contingency changes). The model increases the learning rate when volatility is high and decreasing it when outcomes are noisy. A related meta-learning account showed that serotonergic neurons adjust learning speed according to expected and unexpected uncertainty (10), emphasizing neuromodulatory control over learning dynamics. In our framework, volatility is estimated using an incompatibility term that captures whether the direction of choice updating diverges from that suggested by recent outcomes. Probabilistic reversal (PRL) behavior has also been modeled using inference-based accounts that assume explicit representation of hidden task states – with animals updating their belief that there is a high- and a low-reward probability lever (35, 36). Here we instead adopted an adaptive value-based account in which learning rates vary as a function of recent outcome patterns, capturing uncertainty effects without invoking state inference. Yet, our approach bears a conceptual proximity with inference: our volatility estimate is derived from the incompatibility measure introduced by Mishchanchuk et al. to simplify inference models (35). However, behavioral differences in these two studies likely reflect differences in training history: inference-based strategies may be promoted by deterministic pre-training in their study, whereas our animals were trained directly under probabilistic contingencies. Consistent with this, how OFC reflects hidden states or reward statistics depends on the training stage in temporal (37) or foraging tasks (38). Overall, these findings support that flexible reversal behavior is best captured by models in which the speed of adaptation (expressed as learning rate in the RL framework) dynamically adjust to two types of uncertainty (stochasticity and volatility).

### The role of the orbitofrontal cortex in adaptive behavior

The OFC has long been linked to behavioral flexibility, particularly in situations where learned action–outcome associations must be updated after contingencies change (26). Across species, OFC lesions produce persistent responding to previously rewarded options, indicating a deficit in updating learned associations rather than in response inhibition. Consistent with this notion, we previously showed that ventrolateral OFC inactivation disrupts goal-directed control specifically after a reversal in outcome identity (27), supporting a role for OFC in revising learned associations. In PRL, OFC activity carries the signals required for feedback-based updating. In rats, OFC neurons encode both the expected value of the chosen action and the subsequent prediction error (PE), with earlier and stronger responses than medial prefrontal regions, consistent with a direct role in adjusting values when outcomes deviate from expectation (39). In mice, two-photon calcium imaging further showed that OFC populations encode difference (ΔQ) or sum (∑Q) in option values, and chosen value on a trial-by-trial basis, directly representing the quantities needed to evaluate and revise choice strategies (32).

Local field potential recordings of the OFC provide convergent evidence for this interpretation. In a closely matched rat PRL task, beta power in the OFC strongly differentiated high- and low-probability options prior to a reversal but collapsed immediately after contingencies switched, when rats had to re-accumulate reward evidence (40). Importantly, beta power in lOFC was positively correlated with model-estimated learning rates (40), aligning with our conclusion that OFC inhibition reduced learning rates. Consistent with this interpretation, in a similar rat PRL task, pharmacological OFC inhibition also selectively impaired post-reversal adjustment and decreased learning-rate parameters without affecting performance during stable periods (29). Disrupting OFC function also reduces adaptation to feedback on low-probability option, which is critical for adaptation in probabilistic environments (41). In mice, CaMKII-dependent plasticity in OFC supports trial-by-trial RL, and transient silencing the OFC abolishes the influence of recent outcome history on choice, indicating that OFC activity is required for using recent feedback into behavioral adjustment (32). However, not all studies report equivalent OFC involvement. For example, Jeong et al. (42) used a mouse two-option probabilistic spatial bandit task -conceptually similar to Sul et al. (39) in rats- and found minimal behavioral effects of OFC perturbation. These discrepancies likely reflect task demands. For instance, a study in Macaques (43) reported behavioral deficits in a multi-outcome reversal task similar to those observed in our study, whereas no impairment was observed in a simpler, deterministic two-option reversal version of the same task - a pattern also reported in rats (41). This suggests that OFC contributions become especially critical when adapting behavior requires integrating uncertainty and rebuilding value estimates after contingency changes.

As noted above, the volatility structure of our PRL task is performance-dependent, leading to relatively stable reward periods before each contingency change. Prior work indicates that OFC contribution to reversal learning depends on such stability: when reversals are relatively infrequent, allowing animals to accumulate evidence about the current action–outcome relationship, OFC plays a key role in adjusting behavior when that relationship changes. However, when reversals occur very frequently (≈5 trials), outcome expectancies cannot accumulate, and OFC involvement is reduced or can even hinder performance (44, 45). The stable pre-reversal periods in our task likely enhance reliance on OFC mechanisms for adjusting behavior when expectations are violated.

In addition, multi-option probabilistic reversal tasks demonstrate similar patterns of OFC involvement. In a three-armed bandit task, disrupting OFC→nucleus accumbens core projections reduced learning from negative feedback and slowed post-reversal adaptation (28), indicating that OFC supports value updating even when multiple competing action values must be evaluated. This suggests that the role of OFC in reversal learning is not limited to two-option settings, but extends to situations requiring the resolution of several alternative value representations. These observations also suggest that our own post-reversal effects may likewise depend, at least in part, on OFC→striatal projections.

### Noradrenergic modulation of the orbitofrontal cortex for adaptive behavior

Yu and Dayan (12) proposed that LC–NA signals unexpected uncertainty, increasing the learning rate when outcome variability exceeds expectations. Sales et al. (13) similarly framed LC–NA as a state–prediction error that dynamically adjusts learning speed, enabling shifts between stable exploitation and flexible updating when contingencies change. Our results are in line with recent human studies that have investigated the role of NE in modulating learning under conditions of uncertainty. In predictive-inference tasks, Nassar et al. (19) have shown that pupil dilation, often seen as a proxy for LC–NA activity, transiently increases following changepoints and covaries with trial-by-trial adjustments in learning rates. Pharmacological manipulation of NA modulates learning rate in a baseline-dependent manner, with the strongest effects following subtle or ambiguous contingency changes (46, 47). In aversive learning, participants normally increase their learning rate in volatile environments (6), an adjustment tracked by pupil-linked noradrenergic responses; and individuals with higher anxiety showed reduced volatility-based learning-rate adaptation alongside a blunted pupil response to volatility. Catecholamine enhancement with methylphenidate also increased the ability to scale learning rates with volatility in a probabilistic bandit task (48), whereas β-adrenergic blockade impaired volatility-driven learning-rate adjustments and reduced pupil–volatility coupling (49). Overall, these findings suggest a causal role for noradrenergic signaling in regulating learning speed when contingencies change.

Recent rodent work similarly supports a role for LC–NA in adaptive updating. In mice, baseline LC firing predicts and drives faster rule updating in a Go/No-Go reversal task (50). In rats, noradrenergic input specifically to ventrolateral OFC is required for updating reversed specific action–outcome associations, whereas orbitofrontal dopamine and LC→mPFC noradrenergic pathways do not produce comparable effects (51).

Extending these findings, we observed that NA release in the OFC increased specifically at moments requiring behavioral updating, likely providing an estimate of trial-by-trial volatility as in our adaptive learning model. Chemogenetic inhibition of LC→OFC projections reproduced the behavioral deficits predicted by disrupting volatility-driven learning-rate modulation, selectively impairing post-reversal adaptation in probabilistic conditions while sparing performance under deterministic contingencies.

This convergence across model predictions, neural dynamics, and causal perturbation indicates that noradrenergic LC projections to OFC support the dynamic regulation of learning speed when environmental contingencies change.

## Methods

### SUBJECTS AND HOUSING CONDITIONS

A total of 59 Long–Evans rats, aged 2–3 months, were acquired from the Centre d’Elevage Janvier (France). Animals were housed in pairs with ad libitum access to water and standard lab chow, then put on food restriction 2 days before the start of behavioral experiments, and maintained at ∼90% of their ad libitum feeding weight. Rats were handled daily for 4 days before the beginning of the experiments. The facility was maintained at 21 ± 1°C on a 12h light/dark cycle (lights on at 8:00 A.M.). Environmental enrichment was provided by tinted polycarbonate tubing elements, in accordance with current French (Council directive 2013-118, February 1, 2013) and European (directive 2010-63, September 22, 2010, European Community) laws and policies regarding animal experimentation. The experiments received approval from the ethics and animal experimentation committee of the French Ministry of Higher Education and Innovation (reference number of the project: APAFIS#27928-2020110918011853 v2).

### VIRAL VECTORS

For chemogenetic inhibition of the orbitofrontal cortex, we used an adeno-associated viral (AAV) vector encoding the inhibitory hM4Di DREADD (designer receptor exclusively activated by designer drugs). The AAV8-CaMKII-hM4D(Gi)-mCherry virus was obtained from the Viral Vector Production Unit at Universitat Autònoma de Barcelona, Spain. This construct was derived from the Addgene plasmid #50477 (pAAV-CaMKIIa-hM4D(Gi)-mCherry) and was delivered at a titer of 3.5 × 10¹² GC/ml, as previously validated (Parkes et al., 2018). To selectively inhibit noradrenergic projections to the orbitofrontal cortex, we used a retrograde canine adenovirus type 2 (CAV-2) vector under the control of the PRS promoter (a noradrenergic-specific cis-regulatory element from the human dopamine β-hydroxylase gene). The CAV-2-PRS-HA-hM4D(Gi)-hSyn-mCherry viruses were provided by Marina Lavigne and Eric J. Kremer (Institut Génétique Moléculaire de Montpellier, CNRS UMR 5535, France) and used at a concentration of 3.5 × 10¹² VP/ml, as previously validated in (51, 52). For fiber photometry recordings, we employed the genetically encoded noradrenergic sensor GRAB_NE1m to monitor real-time noradrenaline release in the target region. The AAV9-hSyn-GRAB_NE1m virus was obtained from Addgene (#12308-AAV9) and was originally developed and validated by Yulong Li’s laboratory (Peking University, Beijing, China). The virus was used at an undiluted working titer of 1 × 10¹³ vg/ml (53).

### STEREOTAXIC SURGERIES

For all experiments, rats were anaesthetized with 5% inhalant isoflurane gas with oxygen and placed in a stereotaxic frame with atraumatic ear bars (Kopf Instruments) in a flat skull position. Anaesthesia was maintained with 2% isoflurane and complemented with a subcutaneous injection of ropivacaïne and lidocaïne (a bolus of 0.2 mL 2mg/mL) at the incision site and following disinfection using Vetedine 10% and 5%. All viral vectors were injected using a microinjector (UMP3 UltraMicroPump II with Micro4 Controller, World Precision Instruments) connected to a 10µl Hamilton syringe. For OFC inactivation and LC to OFC inactivation experiments, each animal received two injections per hemisphere of either AAV8-CaMKII-hM4D(Gi)-mCherry or CaV2-PRS-HA-hM4D(Gi)-hSyn-mCherry. Each injection consisted of 1µl infused at a rate of 200nl/min. After each injection, the syringe remained in place for an additional 5min before being removed. The coordinates used for the OFC (VO + LO) were the following (in mm from Bregma): AP +3.7, ML ± 2.0, DV −5.0 and AP +3.2, ML ± 2.8, DV −5.2 for both AAV8-CaMKII-hM4D(Gi)-mCherry and CAV-2-PRS-HA-hM4D(Gi)-hSyn-mCherry injections and AP +3.45, ML ± 2.4, DV −5.1 for AAV9-hSyn-GRABNE1m. During surgery, a heating pad was placed under each rat to maintain its body temperature. For photometry experiments, a blunt-ended fiber optic cannula (400 µm, Aquineuro) was carefully implanted 200 µm above the injection site and secured using highly adhesive dental cement (Super-Bond, C&B) in combination with four stainless steel jeweler’s screws. Following surgery, rats received a subcutaneous injection of the non-steroidal anti-inflammatory drug meloxicam (2 mg/kg) and were individually housed in warmed cages with facilitated access to food and water for 2 hours post-surgery. A recovery period of two weeks was allowed to ensure proper transgene expression before behavioral training. Injection sites were later confirmed histologically upon completion of the experiments.

### CHEMOGENETICS

The DREADDs agonist deschloroclozapine (DCZ, MedChemExpress) was dissolved in dimethyl sulfoxide (DMSO) at a concentration of 50 mg/ml and stored at −80°C as a stock solution. On the day of use, the stock was diluted in sterile saline to a final concentration of 0.1 mg/kg and administered intraperitoneally (10 ml/kg, i.p.) 35 minutes before behavioral testing, following validation in a previous study (51). The vehicle (Veh) solution consisted of 0.2% DMSO in sterile saline. All injectable solutions were freshly prepared and handled under low-light conditions.

### HISTOLOGY

At the end of behavioral experiments, rats were sacrificed with an overdose of sodium pentobarbital (Exagon Euthasol) following an i.p. injection of xylazine (20 mg/kg) and perfused transcardially with 60ml of saline followed by 260ml of 4% paraformaldehyde (PFA) in 0.1M phosphate buffer (PB). Brains were removed and postfixed in the same PFA 4% solution overnight and then transferred to a 0.1M PB solution. Subsequently, 40µm coronal sections were cut using a VT1200S Vibratome (Leica Microsystems). Every fourth section was collected to form a series. Immunofluorescence staining was performed for mCherry (AAV8-CaMKII-hM4D(Gi)-mCherry) and GFP (AAV9-hSyn-GRABNE1m) proteins. For red fluorescent proteins, free-floating sections were first rinsed in 0.1M phosphate-buffered saline (PBS; 4 × 5min) and then transferred to PBS containing 0.3% Triton X-100 (PBST) for further rinsing (3 × 5min), before being incubated in blocking solution (PBST containing 3% goat serum) for 1h at room temperature; sections were then incubated for 24h at room temperature with a primary antibody (rabbit polyclonal anti-RFP, 1/1,000) diluted in blocking solution; after washes (4 × 5min) in PBST, sections were incubated for 2h at room temperature with the secondary antibody (goat polyclonal anti-rabbit TRITC-conjugated, 1/200) diluted in PBST; following rinses in PBS (4 × 5min), sections were collected on gelatin-coated slides using 0.05M PB, before being mounted and coverslipped using Fluoroshield with DAPI mounting medium. DAB staining was performed for mCherry to confirm CAV-2-PRS-HA-hM4Di-hSyn-mCherry injection sites: free-floating sections were rinsed in PBST (4 × 5min) before being incubated in an H2O2 (0.5%) in PBST solution for 30min at room temperature; sections were rinsed again in PBST (4 × 5min) before being incubated in the blocking solution; following blocking, they were incubated overnight at room temperature with the primary antibody (rabbit polyclonal anti-RFP, 1/2,500) in blocking solution; after washing in PBST (4 × 5min), sections were placed with the secondary antibody (biotinylated goat anti-rabbit, 1/1,000) diluted in PBST for 2h at room temperature; following washes (4 × 5min) in PBST, they were then incubated with the avidin–biotin–peroxydase complex (1/500 in PBST) for 90min in a dark environment at room temperature. For the final DAB staining, H202 was added to the solution (10mg tablet dissolved in 50ml of 0.1M Tris buffer) just before incubation of 3–4min; sections were then rinsed with 0.05M Tris buffer (2 × 5min) and 0.05M PB (2 × 5min), before mounting in 0.05 M PB and coverslipping using a Fluoroshield medium.

### BEHAVIORAL APPARATUS

Behavioral training and testing were conducted in eight identical operant chambers (40 cm width × 30 cm depth × 35 cm height, Imetronic), each housed within a sound- and light-resistant wooden enclosure (74 × 46 × 50 cm). Each chamber featured a pellet dispenser that delivered 45 mg grain-based pellets (Bio-Serv) into a food magazine upon activation. Two retractable levers were positioned on either side of the magazine. A ventilation fan provided a constant background noise of 55 dB, and four ceiling-mounted LEDs illuminated the chamber. Experimental events were controlled and recorded using a computer running POLY software (Imetronic). Prior to testing, rats underwent two daily habituation sessions, during which they received 30 grain pellets in the magazine at pseudorandom intervals averaging 60 seconds.

### PRE-TRAINING

#### Initial Lever Press Training

Rats were initially trained to press each lever under a fixed-ratio 1 (FR1) schedule of reinforcement. Levers were inserted alternatively into the cage for 10 minutes or until 20 lever presses. Each lever was presented twice per session, and a single grain pellet was delivered upon each press. Training sessions continued daily until rats reached a criterion of 80 presses across two consecutive days.

#### Simplified Task Training

Before proceeding to the full task, rats underwent training on a simplified version of the task designed to familiarize them with probabilistic reward delivery and equalize the value associated with each lever.

Each session consisted of 90 trials. At the start of each trial, both levers were retracted, and the house light was turned off. Every 20 seconds, a new trial was initiated with the illumination of the house light and the insertion of one of the levers. The order of insertion of the two levers was pseudo-random. Rats were given a 10-second window to press the lever. Failure to press within the allotted time resulted in the retraction of the lever and the house light being turned off, which was recorded as an omission. A press within the 10-second window led to a 50% probability of a grain pellet reward. Training sessions were conducted daily, until rats could complete more than 80 trials for two consecutive days to meet the criterion for progression to the full task.

### BEHAVIORAL TASK

#### Reversal learning task

The behavioral procedure was based on previously established paradigms (31, 41, 54). Daily sessions consisted of 200 discrete choice trials. Each trial began with the illumination of the house light for 2 seconds, followed by the simultaneous insertion of both levers into the operant chamber for 10 seconds.

One lever was designated as the “high” lever, associated with a higher probability of reward delivery upon a press. The other lever was designated as the “low” lever, associated with a lower probability of reward delivery. Upon a lever press, both levers were immediately retracted, and any rewards were dispensed into the magazine. If the rat failed to press either lever within the 10-second window, the levers were retracted, and the house light was turned off until the initiation of the next trial. Across the session, each time the high lever was selected for eight consecutive trials, the contingencies were reversed so that the high lever now became the one that provided a lower probability of reward (i.e., low lever) and vice versa.

For analyses in Fig. 1, we defined training stages based on session number under the initial 80/20 schedule. Naïve rats corresponded to Sessions 1–2 of training, and intermediate rats to Sessions 9–10, reflecting early and late acquisition stages before reaching expert performance.

After reaching expert performance on the 80/20 schedule, rats were successively exposed to different reward probability pairs (70/30, 90/10, and 100/0; Fig. 3). For each contingency, rats performed 8 daily sessions; behavioral analyses were restricted to the last 4 sessions of each contingency. Between each change in contingency, animals were re-trained on the original 80/20 schedule to maintain a common baseline. Thus, for comparisons across contingencies, we used 4 sessions per condition, including the 80/20 schedule.

For photometry recordings, each trial began with illumination of the house light for 1s, followed by simultaneous insertion of both levers into the operant chamber for 8 seconds, within a session lasting 30 minutes.

For chemogenetic experiments (Fig. 2 and 5), rats were first trained to stable performance on the probabilistic 80/20 reversal-learning task. Once performance stabilized, each animal underwent vehicle and DCZ test sessions under these probabilistic contingencies (two sessions of each type, in a counterbalanced order). For the LC→OFC experiment, following these probabilistic tests, rats were switched to a deterministic schedule (p(high) = 1.0, p(low) = 0.0) for four daily training sessions. After this deterministic training period, animals again underwent vehicle and DCZ test sessions (two sessions of each type, counterbalanced).

### DATA ANALYSIS

*Win–stay / lose–shift behavior* was measured to analyse feedback sensitivity (54). The win–stay ratio was calculated as the proportion of trials in which animals repeated a choice after it was rewarded. Conversely, the lose–shift ratio was calculated as the proportion of trials in which animals switched after a non-rewarded trial. Ratios were computed separately for correct and incorrect lever choices. In addition to WS/LS analyses, we quantified the proportion of switching choices as a function of whether the previous choice was the high-probability or low-probability lever and whether that choice was rewarded or not. This produced four conditions: (i) high-probability rewarded, (ii) high-probability unrewarded, (iii) low-probability rewarded, and (iv) low-probability unrewarded. This analysis allowed us to assess how outcome history and lever value jointly influenced the likelihood of switching behavior.

*Behavior* was also quantified by computing the number of reversals completed per session and normalizing this value to the number of reversals per 100 completed trials. This controls for variation in trial omissions, ensuring that changes in reversal performance were not driven simply by differences in total trials completed.

*Perseverative errors* were defined as the number of consecutive incorrect responses following a reversal of reinforcement contingencies, up to the first correct response. Errors committed after the first correct choice were not counted as perseverative. To equate across sessions in which animals completed a different number of reversals, perseverative errors were averaged across the minimum number of reversals completed under both conditions for each animal.

Proportion of high-reward probability lever-presses around reversals.

*Reversal performance* was quantified by aligning trials to the reversal point (t) and calculating the proportion of high-reward probability lever-presses on each subsequent trial. Choices were averaged across reversals within each session and then across animals, yielding trial-by-trial adaptation curves (t+1, t+2, … t+8) that reflect how quickly behavior adjusted after the contingency change.

For summary analyses, we defined two periods:

- Post-reversal period: the first eight trials following each reversal.
- Stable period: all trials outside of this post-reversal window.

For each animal and condition, we calculated the mean proportion of choices to the high-value lever separately for these two periods. This allowed us to compare adaptation immediately after reversals to performance during stable phases of the task.

*Lever-press latencies* were computed on a trial-by-trial basis as the time elapsed between lever insertion onset and the subsequent lever press. Trials without a press (omissions) were excluded from latency analyses.

*Choice–outcome history* was quantified using a logistic regression approach adapted from Parker et al. (2016), with custom MATLAB scripts. For each contingency condition, data were grouped by rat and session. We then fit separate generalized linear models (GLMs, binomial distribution, logit link) for rewarded and unrewarded outcomes across the four previous trials (lags t−1 to t−4). On each trial, the dependent variable was whether the animal repeated its previous choice (stay = 1, switch = 0). Predictor variables encoded whether the choice on a past trial was rewarded or unrewarded, and the side pressed (left = –1, right = +1). Thus, regression coefficients capture the influence of prior rewarded presses (reward predictor) or unrewarded presses (unreward predictor) on the odds of repeating the same choice. Positive coefficients indicate a bias to repeat after the corresponding outcome, while negative coefficients indicate a bias to switch. Regression models were fitted using the fitglm function in MATLAB. For each rat and contingency, coefficients were extracted at each lag (t–1 to t–4) and averaged across animals. Group means and standard errors were then computed for reward and non-reward predictors separately, providing a measure of how past outcomes shaped ongoing choice behavior.

*Choice–outcome history* was further quantified using conditional switching to assess how recent history influenced current decisions. We used a coding scheme (55) that jointly represents the previous choice and its outcome. In this scheme, the direction of a choice is represented by a letter, with the same letter indicating a repeated choice and a different letter indicating a switch between trials t-2 and t-1. The outcome of each choice is encoded using case, where uppercase denotes a rewarded trial and lowercase represents an unrewarded trial. For example, the sequence Aa indicates that the choice at trial t-1 was unrewarded, while the choice at trial t-2 was rewarded and made on the same lever. Conversely, aB signifies that at trial t-1, the choice was rewarded after switching from the other lever at t-2, which had resulted in no reward. This coding system was used to construct sequences representing prior choice-outcome histories, allowing us to analyze their impact on subsequent decision-making.

### FIBER PHOTOMETRY

Fiber photometry recordings were performed using a multichannel fiber photometry system (R811, RWD Life Science). Excitation light sources at 405 nm and 465 nm were used to measure the reference and GRAB_NE1m fluorescence signal, respectively. Optical fiber patch cords (400μm, AquiNeuro) were connected to a rotary joint and photobleached for at least 2 hours before use. All equipment was supplied and installed by AquiNeuro. Light intensity at the tip of the patch cord was measured before each recording session and maintained at 50 μW for 405 nm and 100 μW for 465 nm. Each recording lasted approximately 32 minutes, including 1 minute before and 1 minute after the behavioral exercise. TTL signals (50 ms pulses) were sent from the operant chamber to the fiber photometry system to mark reward deliveries and lever presses. These markers were timestamped in the fiber photometry recordings using a synchronization algorithm and saved as CSV files. The CSV files were then processed using two in-house custom-made software programs, which enabled analysis of experimenter-defined events and intervals. The fluorescence ratio signal (z-score = [(DF/F – mean session) / stdev session]) was analyzed. For each subject, the average z-scores within the behavioral window of interest were used for statistical analysis. Photometry signals were aligned to lever press and analyzed in predefined epochs relative to lever press: Pre-press (–2.0 to –0.1s), Outcome (+0.5 to +2.0s), and Post-outcome (+5.0 to +7.0s). These windows were selected based on the characteristic temporal dynamics of GRAB_NE1m responses in our task, capturing preparatory activity, rapid outcome responses, and delayed post-feedback activity, respectively.

### COMPUTATIONAL MODELING

We used variants of reinforcement-learning models (7) as generative models of rats behavior in the probabilistic reversal task. All models determined the probability P_i_(t) of choosing the next lever i = {L,R}, as a function (the choice rule) of a decision variable (the value of each lever V_L_ and V_R_). The choice rule was the softmax rule, given by (for instance, for i=L):

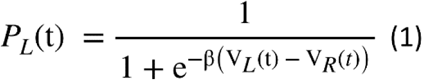

With the explore/exploit parameter β (agents with large β will favor the lever with the largest value more often, agents with low β will tend to choose equally both levers whatever their respective values). The value of a lever was updated using delta rules (Rescorla-Wagner and variants) based on computing reward prediction errors:

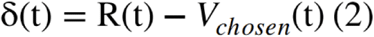

In the simplest formulation, the value update for the chosen lever is:

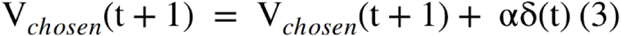

With a learning rate parameter α.

This standard model (with the learning rate being a constant) was fitted to the OFC inhibition data by maximizing the data likelihood (given a sequence of choices on T trials, C(1)…C(T) , data likelihood is the probability of the sequence under the model), with the fmincon function in MATLAB (with the constraints that β is in the ]0,10] range and α in the [0,1] range). This allowed to extract an apparent learning rate for each rat under either vehicle or DCZ conditions.

In the rest of the article (behavior under different levels of stochasticity, model of NA release, and model of LC->OFC inhibition), we compared the properties of the constant learning rate model with an adaptive learning rate model (or “meta-learning” model, (3)), in which the learning rate α(t) is a dynamical variable (instead of a constant parameter) that depends on the running estimates of stochasticity ε(t) and volatility λ(t):

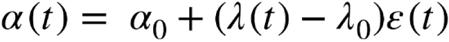

With α_0_ the baseline learning rate and λ_0_ a scaling factor ensuring that the learning rate α(t) is low when ε(t) is high and λ(t) is low (large unsigned errors, low volatility), while α(t) is high when the λ(t) volatility is high even in the presence of large ε(t), as uncertainty can transiently increase when the environment changes (8). We set α_0_ = 0.7 and λ_0_ = 1.3 based on simulations of the ranges of ε(t) and λ(t) so that α(t) would be comprised between 0 and 1. To ensure biological plausibility, we based estimates of ε(t) and λ(t) on reward prediction errors δ(t) only. We used the property that unsigned δ(t) can be used to estimate expected uncertainty (i.e. the expected level of absolute errors; (10, 56)):

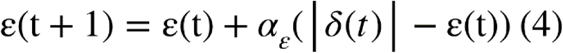

With α_ε_ a constant rate for stochasticity estimation. We kept small (α_ε_ = 0.03) to extract the average stochasticity level outside periods of change. For the volatility estimate λ(t) we considered that animals compute an “incompatibility” term:

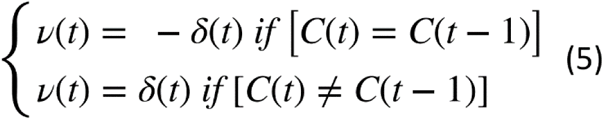

i.e. ν(t) = - δ(t) if rats stayed on their previously chosen option: if they did not change their strategy but obtained less than expected (δ(t) < 0), this signals that the environment has changed and thus ν(t) shall be positive. Conversely, if animals switched from their previous strategy and obtained better than expected, this also signals that the environment has changed. This incompatibility signal is then integrated into a volatility estimate λ(t) by:

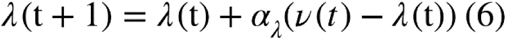

with α_λ_ a constant rate for stochasticity estimation. We used α_λ_ > α_ε_ to extract fluctuations in the environmental statistics that are faster than noise fluctuations (10). We performed model simulations comparison because (i) the constant learning rate model predict opposite directions of effects in perseverative errors or in switching probability conditional on action-outcome sequences compared to adaptive learning rate model – hence providing stronger support for the adaptive learning rate model than quantitative measures of model fitting (33), and (ii) standard model does not include volatility estimates to be compared with the NA release.

### STATISTICAL ANALYSIS

All statistical tests were two-tailed. Data were first tested for normality using the Shapiro–Wilk test, and parametric or non-parametric methods were selected accordingly. Behavioral data were analyzed using paired t-tests, repeated-measures (RM) one-way, two-way, or three-way ANOVAs, or non-parametric equivalents (Friedman test or paired Wilcoxon test) when normality assumptions were violated. Photometry data were analyzed using paired Student’s t-tests, RM one-way or two-way ANOVAs. For analyses requiring multiple comparisons, Sidak’s post hoc correction was applied following significant main or interaction effects. All statistical analyses were performed in GraphPad Prism (version 10), and final figure layouts were assembled in GraphPad and Adobe Illustrator CC.

## RESOURCE AVAILABILITY

Any additional information is available upon request from the corresponding author.

## ACKNOWLEDGEMENTS

This work was supported by the French National Research Agency (CE37-0019 NORAD and CE14-0020 FRONTOFAT to EC), the French government in the framework of the University of Bordeaux’s IdEx “Investments for the Future” program/GPR BRAIN_2030, the “Fondation pour la Recherche sur le Cerveau” (C20492 to EC). HP was awarded a PhD fellowship from the Bordeaux Neurocampus Graduate program and the Iresp (24Iresp017IT07). AP was supported by the Brain & Behavior Research Foundation (Young Investigator Grant 31540). The authors wish to thank: Eric J. Kremer and Marina Lavigne for the conceptualization and generous donation of all CAV2-PRS constructs; AquiNeuro for helping with the fiber photometry set-up; Angélique Faugère for helping with immunohistochemistry and Yoan Salafranque for his expert animal care.

## AUTHOR CONTRIBUTIONS

Conceptualization, HP, JN and EC; methodology, HP, CC, AP, AM, JN, EC; software, AM; formal analysis, HP, CC, JN, AM; investigation, HP, AP, CC; writing – original draft, HP, JN, EC; writing – review & editing, HP, CC, AP, AM, JN and EC; visualization, HP; supervision and funding acquisition, JN and EC.

## Supplementary figures

**Supplementary figure 1:**
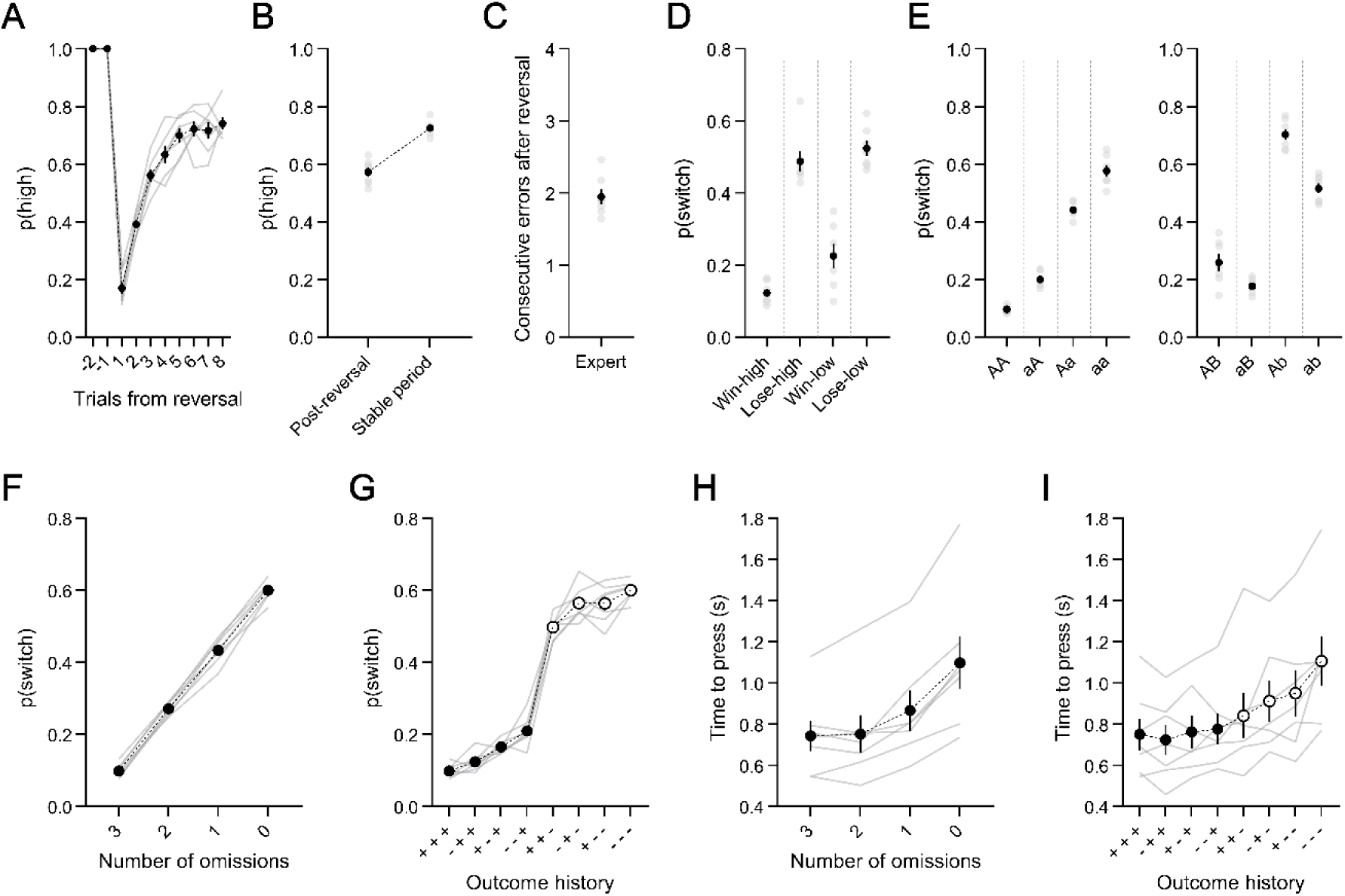
Behavioral performance of OFC-GRABNE1m rats in the probabilistic reversal learning task. (A) Proportion of high-reward probability lever-presses across trials following contingency reversals in expert OFC-GRABNE1m expert rats. (B) Proportion of high-reward probability lever-presses during post-reversal and stable periods. (C) Mean number of consecutive errors following reversals. (D) Conditional probability of switching after rewarded (win) or unrewarded (lose) trials on the high- or low-probability lever. (E) Conditional switching probabilities based on action–outcome sequences in the two preceding trials. The left panel shows sequences where the same lever was chosen (AA, aA, Aa, aa); the right panel shows sequences where different levers were chosen (AB, aB, Ab, ab). (F) Probability of switching as a function of the number of unrewarded outcomes within the last three trials. (G) Conditional switching probabilities decomposed by all possible combinations of outcomes over the preceding three trials (t₋₃, t₋₂, t₋₁; “+” rewarded, “–” unrewarded). (H–I) Lever-press latencies analyzed in the same manner as (F–G), showing reaction times as a function of recent omission count (H) and full outcome history (I).Data are presented as mean ± SEM; each gray line or dot represents one rat.

**Supplementary figure 2:**
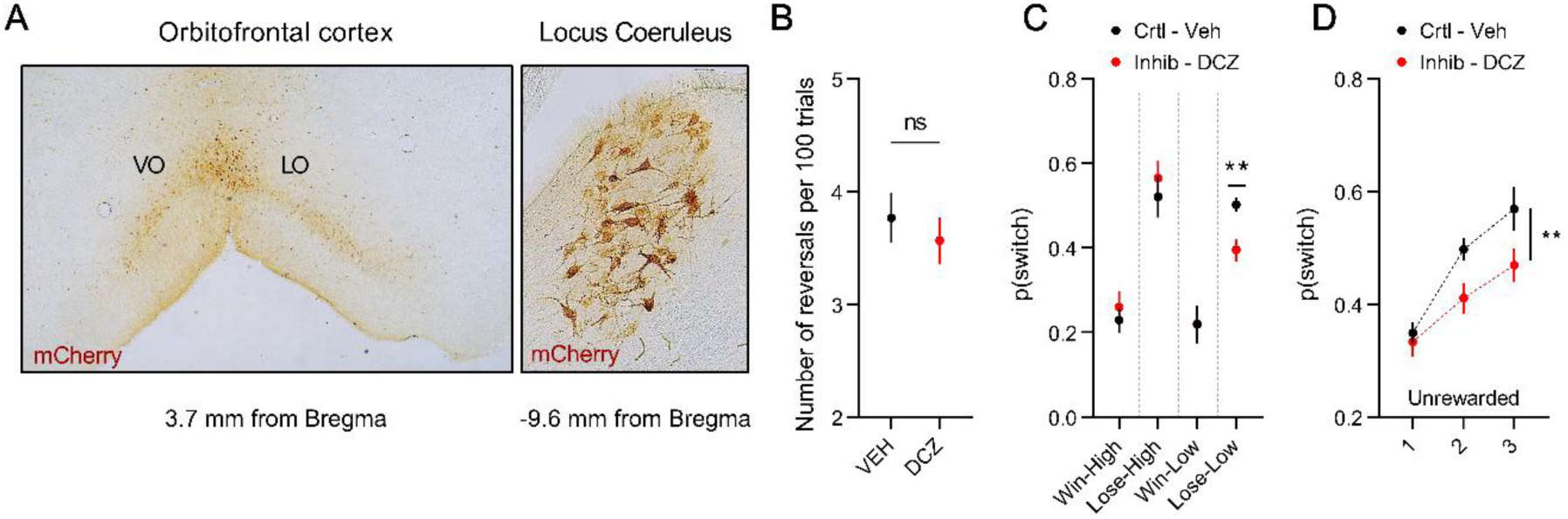
Expression of the LC→OFC chemogenetic construct and behavioral effects of inhibition. (A) Representative images showing the viral injection site in the orbitofrontal cortex (OFC; left) and retrogradely labeled mCherry-positive neurons in the locus coeruleus (LC; right). (B) Mean number of reversals per 100 trials under vehicle (VEH) and deschloroclozapine (DCZ; 0.1 mg/kg, i.p.) treatment, showing no effect of LC→OFC inhibition on the frequency of contingency reversals (n.s.). (C) Probability of switching levers following rewarded (win), unrewarded (lose), high-probability (high), or low-probability (low) lever presses in the post-reversal phase (p < 0.01). (D) Conditional switching probability following unrewarded outcomes, shown as a function of the total number of rewards obtained on the same lever across the preceding three trials (p < 0.01). Data are presented as mean ± SEM.

**Supplementary figure 3:**
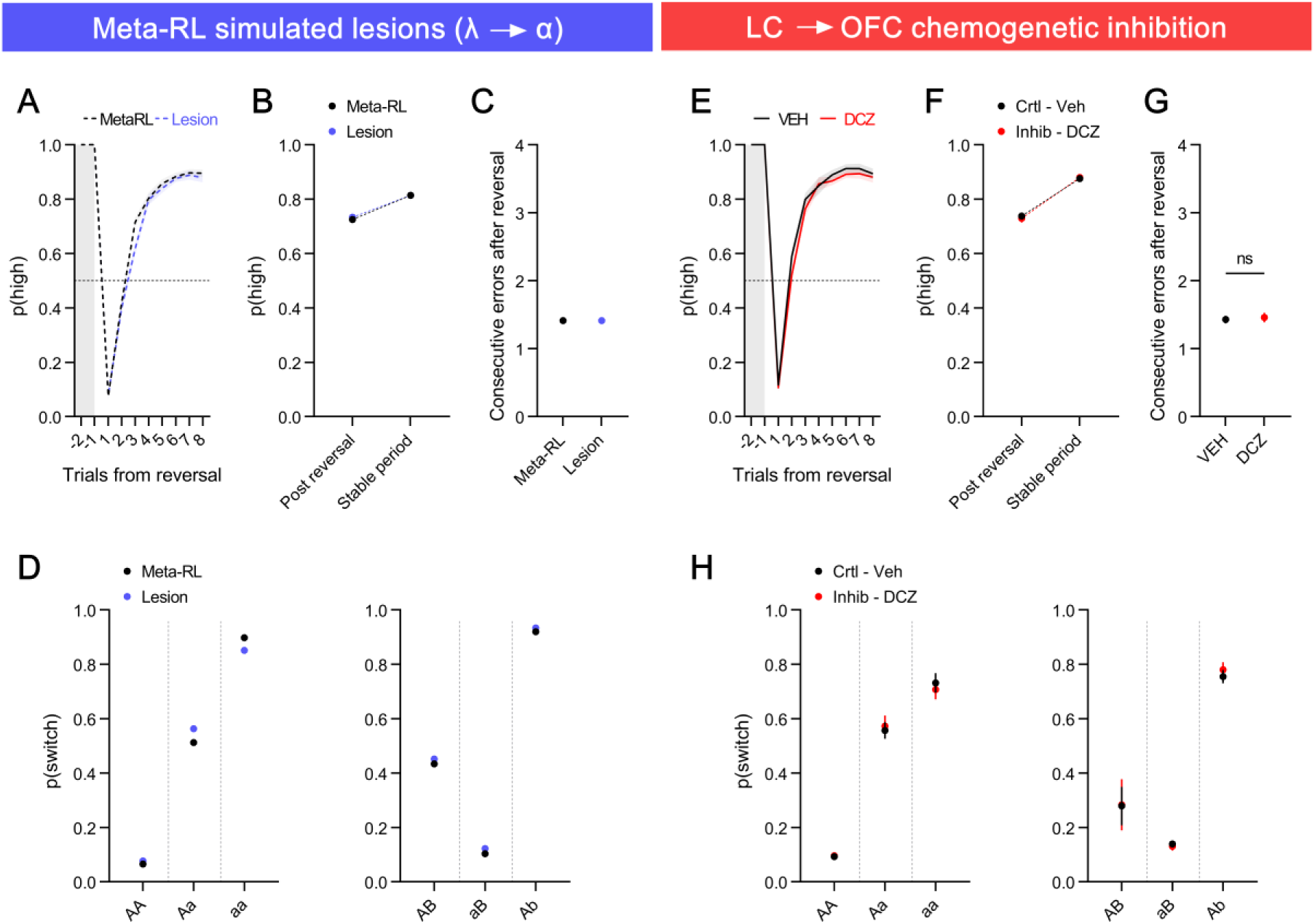
Simulated and chemogenetic LC→OFC inhibition leave adaptive learning intact under deterministic (100/0) contingencies. (A–D) Meta-RL simulations under deterministic conditions. (A) Simulated choice behavior showing the probability of selecting the high-probability lever around reversals for intact (black) and λ→α-lesioned (blue) Meta-RL agents. (B) Proportion of high-reward probability lever-presses during post-reversal and stable periods; no effect of lesion (n.s.). (C) Mean number of consecutive errors after reversals; no difference between intact and lesioned agents (n.s.). (D) Conditional switching probabilities based on the two preceding action–outcome sequences for intact (black) and lesioned (blue) agents, showing no change in switching dynamics. (E–H) In vivo LC→OFC chemogenetic inhibition under deterministic conditions. (E) Behavioral performance across trials following reversals under vehicle (VEH, black) and DCZ (red) treatment. (F) Proportion of high-reward probability lever-presses during post-reversal and stable periods; no treatment effect (n.s.). (G) Mean number of consecutive errors after reversals; no significant difference between VEH and DCZ sessions. (H) Conditional switching probabilities under VEH and DCZ, showing no effect of LC→OFC inhibition on switching behavior. Data are presented as mean ± SEM.

## Statistics table

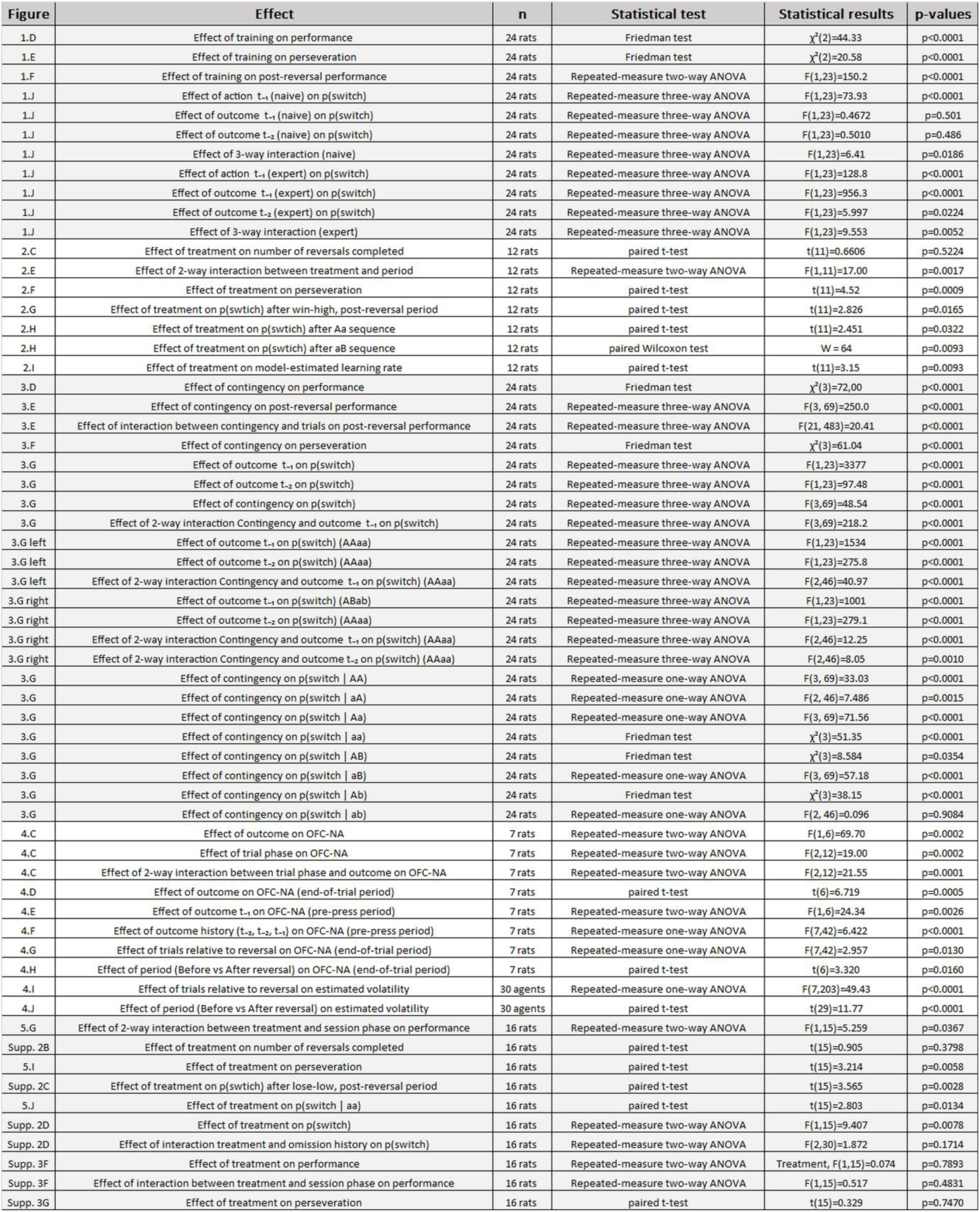

